# Studies on Gene Enhancer with KSHV mini-chromatin

**DOI:** 10.1101/2025.03.24.644916

**Authors:** Tomoki Inagaki, Ashish Kumar, Kang-Hsin Wang, Somayeh Komaki, Jonna M. Espera, Christopher S. A. Bautista, Ken-ichi Nakajima, Chie Izumiya, Yoshihiro Izumiya

**Author notes:** These authors contributed equally to this work (Tomoki Inagaki, Ashish Kumar). Correspondence: Yoshihiro Izumiya DVM, PhD. Address: UCDMC Research III, Room 2200B, 4645 2^nd^ Avenue, Sacramento, CA 95817 E. mail.

## Abstract

Kaposi’s sarcoma-associated herpesvirus (KSHV) genome contains a terminal repeats (TR) sequence. Previous studies demonstrated that KSHV TR functions as a gene enhancer for inducible lytic gene promoters. Gene enhancers anchor bromodomain-containing protein 4 (BRD4) at specific genomic region, where BRD4 interacts flexibly with transcription-related proteins through its intrinsically disordered domain and exerts transcription regulatory function. Here, we generated recombinant KSHV with reduced TR copy numbers and studied BRD4 recruitment and its contributions to the inducible promoter activation. Reducing the TR copy numbers from 21 (TR21) to 5 (TR5) strongly attenuated viral gene expression during *de novo* infection and impaired reactivation. The EF1α promoter encoded in the KSHV BAC backbone also showed reduced promoter activity, suggesting a global attenuation of transcription activity within TR5 latent episomes. Isolation of reactivating cells confirmed that the reduced inducible gene transcription from TR-shortened DNA template and is mediated by decreased efficacies of BRD4 recruitment to viral gene promoters. Separating the reactivating iSLK cell population from non-responders showed that reactivatable iSLK cells harbored larger LANA nuclear bodies (NBs) compared to non-responders. The cells with larger LANA NBs, either due to prior transcription activation or TR copy number, supported KSHV reactivation more efficiently than those with smaller LANA NBs. With auxin-inducible LANA degradation, we confirmed that LANA is responsible for BRD4 occupancies on latent chromatin. Finally, with purified fluorescence-tagged proteins, we demonstrated that BRD4 is required for LANA to form liquid-liquid phase-separated dots. The inclusion of TR DNA fragments further facilitated the formation of larger BRD4-containing LLPS in the presence of LANA, similar to the “cellular enhancer dot” formed by transcription factor-DNA bindings. These results suggest that LANA binding to TR establishes an enhancer domain for infected KSHV episomes. The strength of this enhancer, regulated by TR length or transcription memory, determines the outcome of KSHV replication.

**Importance:** Gene enhancers are genomic domains that regulate frequency and duration of transcription burst at gene promoters, with BRD4 playing a critical role in their enhancer functions. KSHV latent mini-chromosome also contains an enhancer domain made with multiple copies of 801 bp identical repeat DNA fragments, terminal repeats. Here, we utilized manipulable mini-scale chromatins with convenient inducible KSHV reactivation to systematically examine the association between enhancer strength and the outcome of inducible promoter activation. This study illustrated the amount of BRD4 recruitment at the enhancer associated with frequencies of BRD4 distribution to the inducible promoters during KSHV reactivation and, therefore, KSHV lytic replication. Recruitment of BRD4 to the TR is specifically regulated by KSHV latent protein, LANA. KSHV evolves clever enhancer elements designed to be regulated by the KSHV own latent protein, LANA.

## Introduction

The Kaposi’s sarcoma-associated herpesvirus (KSHV) genome consists of a unique coding region flanked by multiple copies of terminal repeat (TR) units [1]. Each TR unit is 801 bp long with high GC content [1, 2], and the number of copies of the TR units varies in KSHV-infected cells [3]. KSHV latency associated nuclear antigen (LANA), encoded by open reading frame (ORF) 73, binds directly to the TR sequences through its C-terminal domain and docks onto host chromosomes via the N-terminal chromatin-binding domain (CBD) [4]. TR sequence contains a DNA replication origin cis-element consisting of two LANA-binding sites (LBS): a higher affinity site (LBS1) and a lower affinity site (LBS2) [3]. While a single copy of the TR containing LBS1 and LBS2 is sufficient for DNA replication through recruitment of the origin replication complexes, at least two TR copies are required for stable episomal maintenance with LANA [3].

In addition to being an Origin of DNA replication, KSHV TR was found to be a transcription regulatory domain for KSHV inducible gene promoters [5, 6]. The studies showed that TR collaborated with KSHV transcription factors, KSHV Replication and transcription activator (K-Rta), and LANA to regulate inducible promoter activity. Since KSHV latent chromatin is circular in infected cells [7, 8], the genomic structure maintains the TR in relatively close proximity to the unique region, which contains an array of inducible viral gene promoters [5]. Similar to cellular enhancer-promoter interactions, the frequency of genomic looping between TR and inducible gene promoters also increases during reactivation [5, 8]. The 3D genomic structure model derived from capture Hi-C data showed that the KSHV 3D genomic structure becomes a compressed doughnut-like shape during reactivation [5, 8]. Here, we examine if multiple TR copies are designed to increase the chance of interactions and enhance the activity of KSHV inducible promoters [9, 10].

Super-enhancers (SEs) are large clusters of gene regulatory elements characterized by high densities of transcription factor binding sites, and these genomic loci often possess active histone modifications, such as acetylation of lysine 27 on histone H3 (H3K27ac) [11, 12]. Bromodomain-containing proteins recognize the H3K27ac-modified histone tail and subsequently tether to the enhancer region [13]. Among multiple bromodomain-containing proteins, Bromodomain-containing protein 4 (BRD4) is the most studied and is known to play a crucial role in the regulation of super-enhancers [14, 15]. Mechanistically, BRD4 facilitates the recruitment of Mediator complex subunit 1 and its interacting proteins on enhancers [16]. Recruitment of transcription factors at the promoters facilitates interaction between enhancers and promoters, partly through components of the mediator complex recruited at promoters. This recruitment increases the local concentration of enzymes by combining transcription related proteins tethered to enhancers and promoters, leading to transcription elongation [16-18]. Such protein interactions are facilitated by liquid-liquid phase separation (LLPS).

LLPS has emerged as a critical biophysical mechanism underpinning gene regulation by concentrating transcriptional machinery and chromatin regulators at specific genomic loci [19]. Transcriptional condensates, such as those formed by BRD4 at SEs, arise from phase-separated assemblies that increase the local concentration of RNA polymerase II, coactivators, and transcription factors, thereby rapidly and effectively amplifying the transcriptional output [20].

Like cellular transcription factors, LANA interacts with BRD4 through a direct protein-protein interaction between the extra-terminal domain of BRD4 and the carboxyl-terminal region of LANA [21]. Previous studies showed that BRD4 is highly enriched at TR and colocalized with RNAPII on KSHV latent chromatins [5]. Using isolated luciferase reporter constructs, the studies showed that the TR possesses gene enhancer function [5]. Ye et al. also showed that LANA and BRD4 co-occupy the host genome and that this genomic region is associated with increased enhancer activity and gene expression in KSHV-infected cells [22].

Here, we investigated the enhancer function of TR by generating recombinant KSHVs with reduced TR copy numbers. We demonstrated that a high TR copy number exerts a stronger enhancer activity and more robustly induces reactivation. This robust reactivation is mediated by the formation of larger LANA NBs, which interact more frequently with inducible promoters but do not increase the frequency of transcription at a specific promoter. Similar to dormant cellular enhancers, we also showed that the TR can be activated by prior transcription activation, forming transcription memory. Finally, we provide evidence that the TR fragment could serve as a platform facilitating LANA-mediated BRD4 containing LLPS *in vitro*.

## Results

### Establishment of recombinant KSHV with varying numbers of TR copies

To assess the transcription function of TR, we first generated mutant KSHV with a reduced number of TR copies using BAC16 recombination [23]. A DNA fragment containing the kanamycin expression cassette and an *I-SceI* restriction enzyme site was introduced into TR fragments with a red recombination system [24]. The number of insertions and position of the kanamycin cassette within TR may vary in each bacterial colony due to multiple identical TR sequence units [2]. Recombination within TR fragments was subsequently induced by the expression of the red recombinase in *E. coli* after induction of *I-SceI* with L-Arabinose (**Figure 1A**). Recombination generated various numbers of TR copies randomly in the KSHV BAC16. After isolating BAC DNA from bacterial colonies, we identified the bacterial clone that harbors 5, 7, 10, or 21 (BAC16 wild type [25]) copies of TR (**Figure 1B**). Each BAC DNA was transfected in iSLK cells, and the stable iSLK cells were established with hygromycin selection (**Figure 1C**). While we attempted multiple times, we failed to establish BAC-stable cells with two copies of TR (TR2). We also noticed that GFP signal intensity was significantly lower in TR deletion mutants, although nearly 100% of iSLK cells showing GFP expression (**Figure 1D, E**).

**Figure 1.**
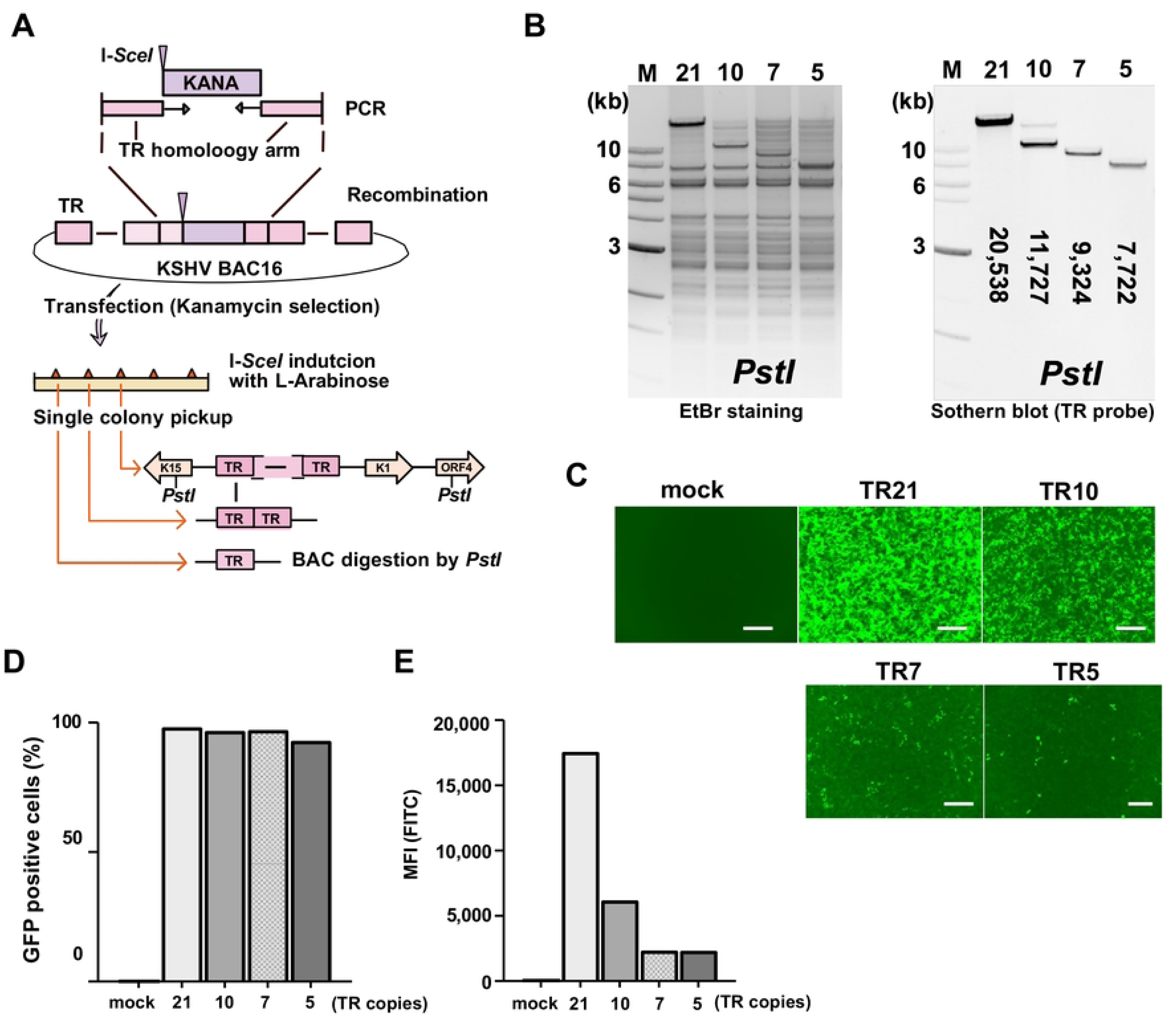
Generation of recombinant KSHV BAC with different numbers of TR copies. (A). Schematic diagram of KSHV BAC16 with different copy numbers of TR. The kanamycin cassette with *I-SceI* recognition sequence, along with the TR homologous sequence, was generated by PCR with pEP-Kan plasmid as a template. This DNA fragment was cloned into KSHV BAC16 by homologous recombination. The kanamycin cassette was deleted by recombination with induction of *I-SceI* in bacteria by incubation with L-arabinose. Correct insertion of the different copy numbers of TR was confirmed by *PstI* digestions. (B). Confirmation of TR copies and BAC integrity. Purified KSHV BAC16 was digested by *PstI* and subjected to electrophoresis. Agarose gel was stained with Ethidium bromide (EtBr) (left panel). The TR-containing fragments were probed with Southern blotting with a fluorescently labelled TR probe (right panel). The numbers in the Southern blotting represent the length (base pair) of the TR fragment after *PstI* digestion. The 1 kb DNA ladder is shown. (C). Fluorescence images of iSLK cells with different copy numbers of TR. KSHV BAC16 with different copy numbers of TR were transfected into iSLK cells. The mock indicates iSLK cells without KSHV BAC16. Scales: 300μm. (D). Proportion of GFP positive cells in iSLK cells with different copy numbers of TR. iSLK cells harboring KSHV BAC16 with different copy numbers of TR were cultured under hygromycin, and the proportion of GFP positive cells was calculated using flow cytometry. E. Mean fluorescence intensity (MFI) of iSLK cells with different copy numbers of TR. MFI in the FITC channel was determined using FlowJo™ v10 software.

### TR is important for viral gene expression

To examine the transcription function of TR in KSHV infection, we first isolated infectious KSHV virions from iSLK cells that were stably transfected with KSHV BAC16 DNAs. The amount of KSHV virion copy was adjusted and infected to iSLK cells at the viral genomic copy/cell ratio of 10:1. We then examined the effect of TR copies in viral gene expression during *de novo* infection. No statistically significant difference was observed in the relative KSHV genome copies in iSLK cells at 24 hours post-infection (**Figure 2A**), indicating that reduced TR copy number does not affect the efficiency of virus infection. Total RNAs were harvested 24 hours post-infection, and RT-qPCR was performed for the latent and inducible viral gene expression. As shown in **Figure 2B**, both KSHV latent and inducible viral genes (*LANA, PAN RNA, vIL-6, and K8.1*) expressed significantly more in KSHV-TR21 infected cells compared to KSHV-TR5 infected cells. Recombinant KSHV-infected cells were then selected with hygromycin, and KSHV-TR21 or KSHV-TR5 latently infected cells were established. Like the BAC transfection, KSHV-TR21 infected iSLK cells showed higher GFP intensity than KSHV-TR5 infected iSLK cells (**Figure 2C**). To examine whether the higher GFP expression is at the transcription level, we next measured GFP transcripts per KSHV genome. The GFP gene is encoded downstream of the EF1α promoter in the KSHV BAC16 backbone and is constitutively expressed in the recombinant KSHV-infected cells [23]. As shown in **Figure 2D**, the GFP was more actively transcribed in the KSHV-TR21 latent chromatin than KSHV-TR5 latent chromatin. We considered that EF1α promoter regulation is independent of KSHV latent protein expression; therefore, this result may suggest that KSHV-TR21 establishes a local nuclear microenvironment that is more favorable for gene transcription.

**Figure 2.**
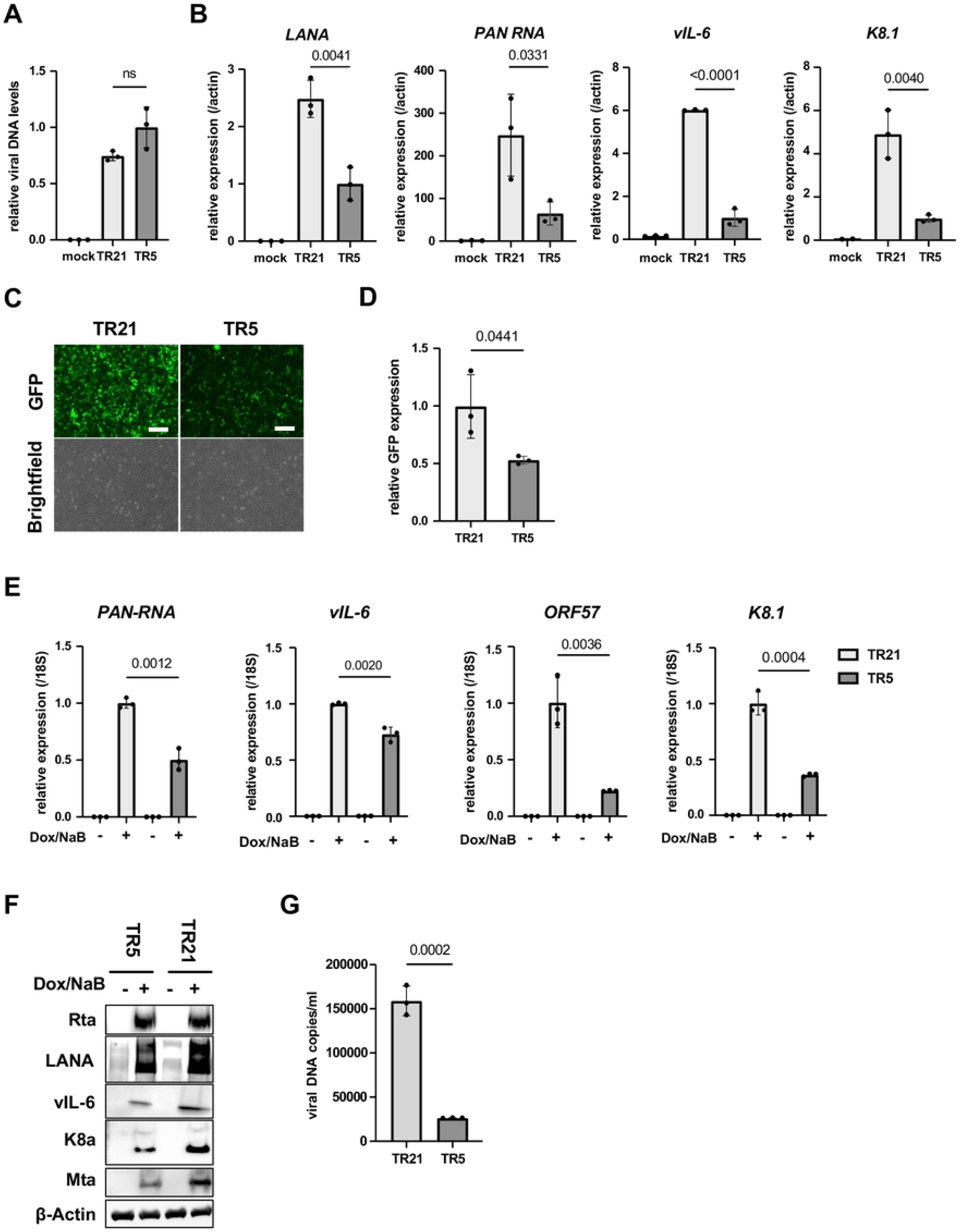
KSHV inducible gene expression is positively regulated in a TR-dependent manner. (A). Relative viral DNA levels in KSHV-infected iSLK cells. iSLK cells were infected with TR5-KSHV or TR21-KSHV for 24 hours and viral DNA levels were determined by quantitative PCR (qPCR). GAPDH expression was used for internal control. Relative viral DNA levels in TR5-KSHV iSLK cells was set as 1. Data was analyzed using a two-sided unpaired Student’s t-test and shown as mean ± SD. (B). KSHV gene expression in iSLK cells during *de novo* infection. Relative RNA levels of the indicated viral genes were determined by Reverse Transcription qPCR (RT-qPCR).18S rRNA expression was used for internal control. Relative gene expression in TR5-KSHV iSLK cells was set as 1. Data was analyzed using two-sided unpaired Student’s t test and shown as mean ± SD. **(C). Fluorescence and brightfield images of iSLK cells infected with TR5-KSHV or TR21-KSHV.** The iSLK cells were infected with the equal amount of TR5-KSHV and TR21-KSHV followed by selection with hygromycin. Scales: 300μm. **(D). Relative GFP expression in iSLK cells infected with TR5-KSHV or TR21-KSHV.** Total RNAs were harvested from iSLK cells infected with the TR5-KSHV or TR21-KSHV. 18S rRNA expression was used for internal control, and relative GFP expression was normalized by the viral genome. Relative GFP expression in TR21-KSHV-infected iSLK cells was set as 1. Data was analyzed using two-sided unpaired Student’s t test and shown as mean ± SD. **(E). Relative KSHV gene expression in iSLK cells infected with TR5-KSHV or TR21-KSHV.** Each iSLK cell was reactivated by sodium butyrate and doxycycline. Total RNAs were purified two days after reactivation. 18S rRNA was used for internal control, and relative gene expression in TR21 iSLK cells after reactivation was set as 1. Data was analyzed using two-sided unpaired Student’s t-test and shown as mean ± SD. **(F). Immunoblotting of KSHV latent protein (LANA) and lytic proteins (Rta, vIL-6, K8α, Mta) in TR5-KSHV or TR21-KSHV-infected iSLK cells.** iSLK cells were reactivated by sodium butyrate and doxycycline and total cell lysates were prepared two days after reactivation and subjected to immunoblotting with specific antibodies. β-actin was used as a loading control. **(G). KSHV virion production after reactivation of TR5-KSHV or TR21-KSHV-infected iSLK cells.** Virions in the supernatant four days after reactivation by sodium butyrate and doxycycline were treated with DNaseI, followed by DNA purification. Results are shown as viral copies per milliliter. ORF6 was used to quantify the viral DNA. The data was analyzed using a two-sided unpaired Student’s t-test and shown as mean ± SD.

Next, we examined the overall efficacies of KSHV reactivation with RT-qPCR. We reactivated KSHV-TR21 and KSHV-TR5 infected iSLK cells with doxycycline and sodium butyrate and examined viral gene expression and KSHV replication. The results showed that the KSHV-TR21 genome was transcribed more robustly after stimulation **(Figure 2E)**, resulting in increased viral protein expression (**Figure 2F**) and virion production in the supernatant (**Figure 2G**). These findings suggest that a higher number of TR copies support gene transcription more efficiently regardless of viral or exogenous genes encoded in the episome.

### BRD4 at the TR region is important for KSHV inducible gene transcription

The results described above led us to hypothesize that cellular and viral proteins at the TR should be important for the KSHV reactivation. This is analogous to the cellular enhancers, which harbor arrays of transcription factor binding sites to maintain transcription-related proteins at the genomic loci [26]. Previous studies showed that LANA interacts with BRD4 [21, 27, 28] and is localized at TR [5, 28]. BRD4 is known to be a localized cellular enhancer [29] and regulates transcription elongation [30]. Accordingly, we focused on whether the shortened TR compromised the recruitment of BRD4 to the viral promoters. Our idea was that the TR recruits BRD4 via sequence-specific LANA DNA binding and that a higher number of TR copies is advantageous for recruiting more BRD4 to the TR through LANA/BRD4 protein interaction; this mechanism established local nuclear environment favorable for transcription activation in the large 3D nuclear space.

First, we examined differences in LANA accumulation on the KSHV episome, using the size of LANA dots as an indicator. An immunofluorescence assay with an anti-LANA monoclonal antibody showed that LANA dots became smaller when we shortened the TR (**Figures 3A and B)**. Proximity ligation assays suggested that larger LANA dots interacted more frequently with BRD4, suggesting that TR21 provides a better platform to recruit BRD4 **(Figures 3C and D)**. We also examined the BRD4 occupancies and histone H3 lysine 27 acetylation (H3K27Ac) in KSHV-TR21 and TR5-infected iSLK cells. As shown in **Figure 3E**, BRD4, and H3K27Ac were more frequently occupied at the TR21 latent chromatin, consistent with the frequency of LANA interactions with BRD4 at LANA nuclear bodies and promoter activity (**Figures 2D and E**).

**Figure 3.**
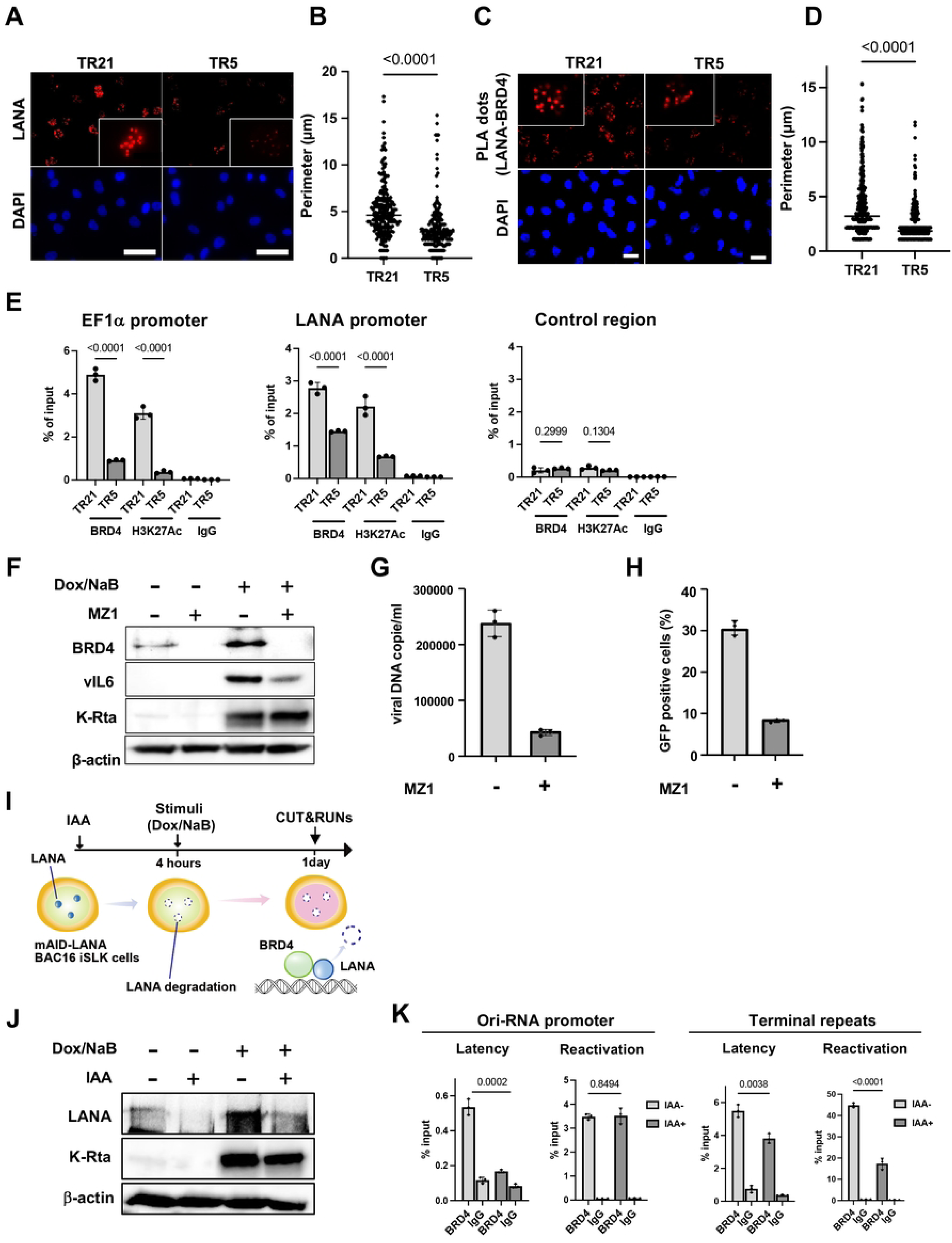
BRD4 regulates KSHV inducible gene expression in the TR dependent manner. **(A). Immunofluorescence image of the LANA protein in TR5-KSHV or TR21-KSHV-infected iSLK cells.** Anti-LANA rat monoclonal antibody and alexa-647 (secondary) goat anti-rat antibody were used for staining LANA in TR5- or TR21-KSHV-infected iSLK cells and Cy5 channel was used for its detection. LANA dots (upper panels) and DAPI (lower panels) are shown. Scales: 50 μm. **(B). Perimeter of each LANA dots**. The size of each LANA dot in TR5- or TR21-KSHV-infected iSLK cells was calculated by imageJ (ver.1.53). Data was analyzed using a two-sided unpaired Student’s t-test and shown as mean ± SD. **(C). Representative images of proximity ligation assay (PLA) for BRD4 and LANA interaction in** TR5- or TR21-KSHV-infected **iSLK cells.** TR5- or TR21-KSHV-infected iSLK cells were subjected to PLA using antibodies against BRD4 (1:100 dilution) and LANA (1:100 dilution). Red dots indicate BRD4-LANA interactions. Scales: 20 μm. **(D). Quantification of PLA dots.** Each PLA dot in TR5- or TR21-KSHV-infected iSLK cells was calculated by imageJ (ver.1.53). **(E). Enrichment of BRD4 and H3K27Ac in** TR5- or TR21-KSHV-infected **iSLK cells.** TR5 or TR21-KSHV-infected iSLK cells were subjected to CUT&RUN analysis with anti-BRD4 (1:100 dilution) or anti-H3K27Ac (1:100 dilution) antibodies to examine their occupancy on the specific promoter regions in the KSHV genome. The control region was the coding region of the ORF23, which has been shown not to accumulate BRD4. Normal Rabbit IgG antibody was used as a negative control. Data was analyzed using a two-sided unpaired Student’s t-test and shown as mean ± SD. **(F). Immunoblotting assay of BRD4 and KSHV proteins.** r.219 iSLK cells were reactivated by doxycycline and sodium butyrate one day after MZ1 (100 nM) treatment. Two days after reactivation, total cell lysates were collected for immunoblotting to detect BRD4 and KSHV proteins (vIL-6 and K-Rta). β-actin was used as a loading control. **(G). Virion production in r.219 iSLK cells with or without MZ1 treatment.** Virions in the supernatant four days after r.219 iSLk cells reactivated by sodium butyrate and doxycycline were treated with DNaseI, followed by DNA purification. MZ1 or DMSO were added one day before reactivation. Results are shown as viral copies per microliter. Data was analyzed using a two-sided unpaired Student’s t-test. **(H). Proportion of GFP positive cells after r.219 KSHV infection.** r.219 iSLK cells were reactivated by sodium butyrate and doxycycline one day after MZ1 or DMSO treatment. Four days after reactivation, an equal amount of supernatant was added to iSLK cells, and the proportion of GFP-positive cells was determined by flow cytometry one day after infection. **(I). Experimental flow of LANA degradation.** mAID-LANA KSHV BAC16 iSLK cells were treated with indole-3-acetic acid (IAA) (5 μM) for four hours, followed by reactivation with sodium butyrate and doxycycline for one day to examine the effect of LANA on BRD4 accumulation on the KSHV genome. **(J). Immunoblotting assay of KSHV gene expression in LANA AID iSLK cells with or without IAA treatment.** One day after reactivation, total cell lysates were collected for immunoblotting to detect KSHV proteins (LANA and K-Rta). β-actin was used as a loading control. **(K). BRD4 occupancy in mAID-LANA KSHV BAC16 iSLK cells.** One day after reactivation. BRD4 enrichment on the Ori-RNA promoter and TR region was determined using CUT&RUN. Normal Rabbit IgG antibody was used as a negative control. Data was analyzed using a two-sided unpaired Student’s t-test and shown as mean ± SD.

To assess the contribution of BRD4 to inducible viral gene transcription, we used MZ1, a selective BRD4 inhibitor [31], and lytic gene expression and viral production were measured. We selected MZ1, a proteolysis targeting chimera (PROTAC) that degrades BRD4, instead of a BET inhibitor. BET inhibitor, acting as an acetylated histone mimetic competitor, would increase the non-chromatin bound BRD4, which may help viral transactivators to form transcriptionally active protein complexes at target gene promoters, and induce KSHV reactivation effectively [32-34]. The immunoblotting and qPCR showed that KSHV inducible protein expression and infectious KSHV virion production were clearly inhibited by the BRD4 degradation (**Figures 3F-3H**).

Next, we examined the role of LANA in BRD4 recruitment on the KSHV latent chromatin. We used mAID-LANA KSHV BAC16 (TR21) iSLK cells [35] to examine whether rapid LANA degradation changes the frequency of BRD4 recruitment. We first incubated the cells with IAA to induce LANA protein degradation for 4 hours and stimulated KSHV reactivation with doxycycline and sodium butyrate (**Figures 3I and J**). The results showed that BRD4 occupancy at the Ori-RNA promoter and TR was significantly decreased by LANA degradation, suggesting that LANA enhances the recruitment of BRD4 to the KSHV episome during latency (**Figure 3K**). As expected, when we triggered KSHV reactivation, BRD4 recruitment was significantly increased. To our surprise, BRD4 recruitment to the Ori-RNA promoter was not affected by LANA degradation after reactivation. On the other hand, BRD4 recruitment to the TR was reduced following LANA degradation. The results indicated that recruitment of BRD4 is differently regulated between TR and lytic inducible promoters. It is important to note that BRD4 recruitment at TR is a magnitude higher than that of the Ori RNA promoter region, and LANA is known to bind TR sequence directly.

### Increased inducible gene expression from a chromatin template with higher TR copies

The experiments above were performed with a cell population in a dish, including non-responder and responding cells with reactivation stimuli. While an increased TR copy number demonstrated increased viral gene expression, it remains unclear whether this outcome is due to an increased number of reactivating cells in the dish, increased viral gene transcription from an episome template, or a combination of both.

To examine whether an increased TR copy number increases the frequency of promoter activation, as an enhancer, we established a new recombinant KSHV, which encodes chicken Bu-1 (cBu-1) extra cell surface domain under the control of the vIL-6 promoter. The vIL-6 coding sequence was then fused with the cBu-1 with the GSG-P2A peptide sequence. The vIL-6 promoter was selected because vIL-6 is highly transcribed among lytic inducible genes during reactivation [36-38] and is one of the direct K-Rta target genes [39, 40]. The GSG-P2A peptide allows two proteins to be translated from the same mRNA by causing the ribosome to fail at making a peptide bond [41-43] (**Figure 4A**). Human cells do not express the chicken protein on their cell surface. Using cBu-1 monoclonal antibody increases the detection of vIL-6 transcribing cells by amplifying signals with fluorescent-labeled secondary antibodies. This strategy identifies and conveniently isolates reactivating living cells. **Figure 4B** demonstrates the co-expression of cBu-1 molecule and vIL-6 protein in a cell. Using protein A/G magnetic beads (**Figure 4C**), we could isolate the vIL-6 transcribing cell population from the dish, which increases the resolution of downstream analyses approximately 10 times by eliminating non-responder cells (**Figures 4D and E**).

**Figure 4.**
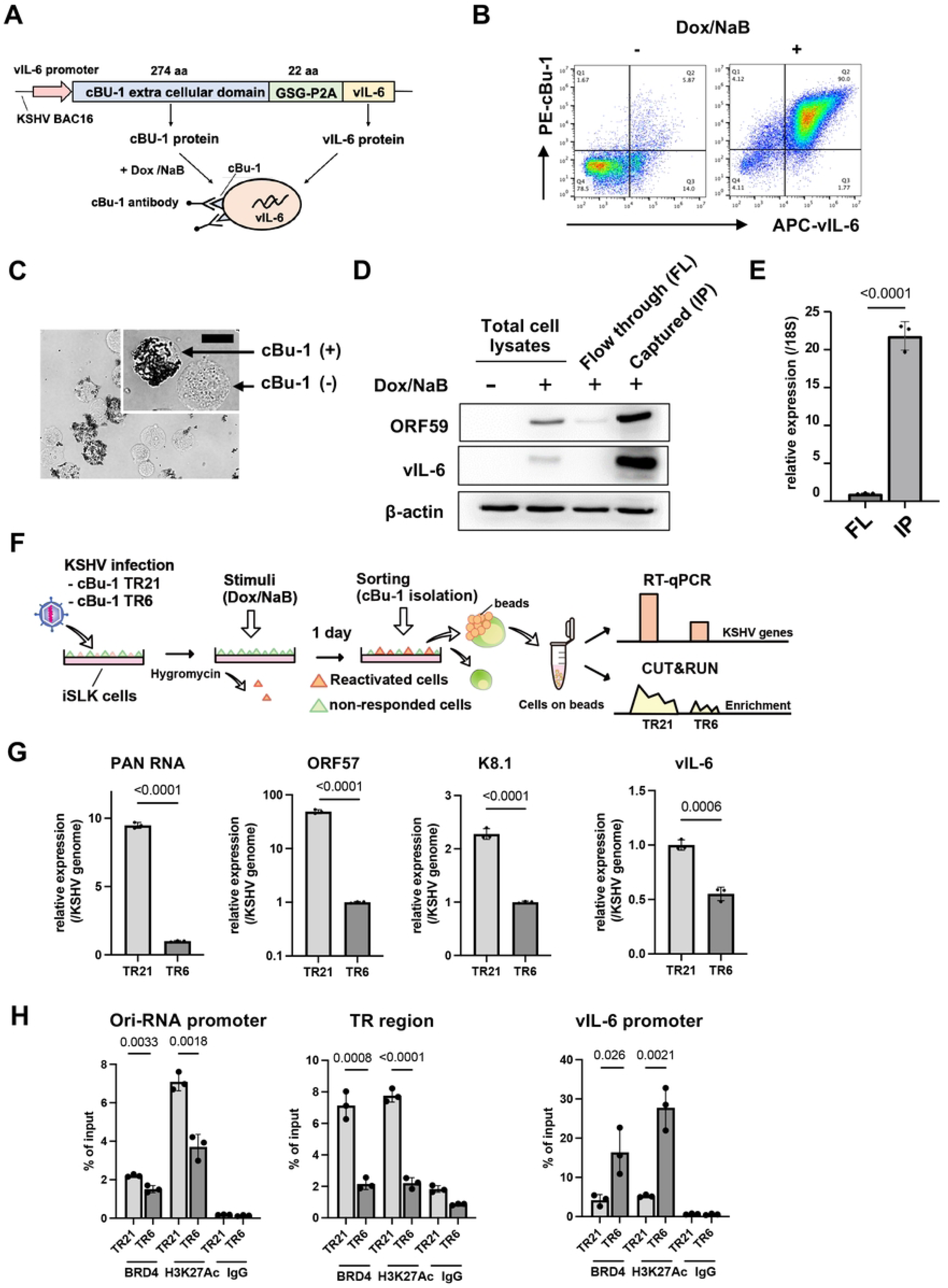
Establishment of cBu-1 BAC16 for the isolation of reactivated cell population. **(A). Schematic model of the isolation of reactivated cells using chicken Bu-1 (cBu-1) BAC16 system.** cBu-1 external domain sequence along with GSG-P2A peptide sequence was inserted as an N-terminal fusion with vIL-6 protein. After reactivation by doxycycline and sodium butyrate, cBu-1 is expressed on the cell surface only in the vIL-6 expressed cells, and anti-cBu-1 antibody can bind to cBu-1 protein for isolation. **(B). Representative FACS profiles of iSLK cells with cBu-1 BAC (cBu-1 iSLK cells)**. Two days after reactivation by sodium butyrate and doxycycline, cells were fixed by 2% paraformaldehyde and permeabilized by 0.1% Triton X-100. Cells were then stained by vIL-6 antibody and cBu-1 antibody for two hours at room temperature. After washing with PBS, cells were incubated with secondary antibodies (APC for vIL-6 and PE for cBu-1). **(C). Brightfield image of cBu-1-expressed cells and non-expressed cells.** Two days after reactivation, cells were incubated with a biotin-conjugated anti-cBu-1 antibody (10 μg/ml) for one hour at room temperature. Cells were washed with PBS twice, followed by incubation with streptavidin magnetic beads for one hour at room temperature. Cells were then observed on the cover glass. Scales: 20 μm. **(D). Immunoblotting of KSHV lytic proteins (ORF59 and vIL-6) in cBu-1 iSLK cells with or without cBu-1 enrichment after reactivation.** cBu-1 iSLK cells were reactivated by sodium butyrate and doxycycline for two days, and cBu-1 expressed cells (IP) were then enriched with cBu1 monoclonal antibody (10 μg/ml). The cell population that did not express cBu-1 was also harvested as flow thorough at the same time (FL). β-actin was used as a loading control. **(E). vIL-6 expression of cBu-1 expressed cells (IP) and non-expressed cells (FL) after isolation by cBu-1 antibody.** 18S rRNA expression was used for internal control. Relative gene expression in FL population was set as 1. Data was analyzed using a two-sided unpaired Student’s t-test and shown as mean ± SD. **(F). Experimental design of the isolation of reactivated cells by cBu-1 system for the following analysis.** cBu-1 virus with different copy numbers of TR (cBu-1 TR21 or cBu-1 TR6) were infected with iSLK cells and cultured with hygromycin. One day after reactivation by sodium butyrate and doxycycline, cBu-1 expressed cells (IP), and cBu-1 non-expressed cells (FL) were sorted by magnetic beads for the following analysis. **(G). Relative KSHV gene expression in iSLK cells after cBu-1 enrichment.** cBu-1 TR21- or cBu-1 TR6-KSHV-infected iSLK cells were reactivated by sodium butyrate and doxycycline. One day after reactivation, reactivated cells were isolated according to cBu-1 expression, and total RNAs were purified to determine the viral gene expression. 18S rRNA was used as an internal standard for normalization, and relative gene expression was normalized by the viral genome. Relative gene expression in cBu-1 TR6-KSHV-infetced iSLK cells was set as 1. Data was analyzed using two-sided unpaired Student’s t test and shown as mean ± SD. **(H). BRD4 and H3K27Ac occupancy in** cBu-1 TR21- or cBu-1 TR6-KSHV-infected **iSLK cells after cBu-1 enrichment.** cBu-1 TR21- or cBu-1 TR6- KSHV-infected iSLK cells were reactivated by sodium butyrate and doxycycline. One day after reactivation, reactivated cells were isolated according to cBu-1 expression, and BRD4 or H3K27Ac enrichment of the Ori-RNA promoter TR, and vIL-6 promoter region was determined. Normal Rabbit IgG antibody was used as a negative control. Data was analyzed using a two-sided unpaired Student’s t-test and shown as mean ± SD.

With the cBu1-vIL-6 recombinant BAC in our hand, we again generated recombinant KSHV with reduced TR copies. While we screened over 50 colonies, we could not find the KSHV BAC possessing TR5, and the closest one was TR6. Accordingly, we used the TR6 clone in the following experiment (cBu-1 TR6-KSHV-infected iSLK cells). As shown in **Figure 4F**, we isolated reactivating cells from cBu-1 TR6 or cBu-1 TR21-KSHV-infected iSLK cells; total RNA was extracted from these cells, which were attached to the magnetic beads. Similarly, CUT&RUNs were performed on the isolated reactivated cell population on the magnetic beads. The relative abundance of BRD4 recruitment per genome copy during active transcription was compared between TR21 and TR6. The robustness of inducible viral genes with stimuli in the reactivating cells was also measured with RT-qPCR. Consistent with the results shown above, TR6-KSHV -infected iSLK cells demonstrated decreased inducible gene expression per genome in the reactivated cells compared to TR21KSHV-infected iSLK cells (**Figure 4G**). Decreased gene expression was accompanied by the decreased frequency or duration of BRD4 recruitment at the inducible promoter (**Figure 4H**). However, when we directly compared the actively transcribing gene (vIL-6) between TR6 and TR21, we found much smaller differences in transcription rates (**Figure 4G**). The recruitment of BRD4 and H3K27Ac modification at the vIL-6 promoter also did not follow a similar trend as other promoters (**Figure 4H, rightmost panel**). The results suggested that increased transcription at PAN RNA or Ori-RNAs in TR21 may be due to a increased frequency of interactions between TR and inducible promoters. Increased interactions likely to lead a greater number of actively transcribing episomes in the infected cell, resulting in an overall increase in inducible viral transcription in the cBu-1(+) cell population.

### Dynamic LANA NBs regulation by transcription activation and the formation of transcription memory

Previous reports showed that KSHV reactivation is a highly heterogeneous process, with only a selected cell population supporting KSHV reactivation in the dish [27, 44, 45]. To understand the underlying heterogenic responses, we isolated iSLK cell populations that could support KSHV reactivation and established two cell populations originating from the same dish (**Figure 5A**). This step allowed us to synchronize the cell condition and compare two stable cell cultures. The isolated cells were cultured for an additional 10 days. The size of LANA nuclear bodies (NBs) (enhancer sizes) was examined, and inducible gene transcription was analyzed using RT-qPCR (promoter activation). Interestingly, the size of LANA NBs was significantly larger in the IP cell population (previously reactivated cell population) than flow through (FL) cells (non-responder with prior stimuli) (**Figures 5B and C**). The number of LANA NBs was comparable between the two cell populations (**Figure 5B**). The RT-qPCR showed that the reactivated cell population (IP) supported inducible gene expression approximately 4-10 times better than the non-responded cell population (FL) (**Figure 3D**). The results suggest that (i) size of LANA NBs are regulated by local transcription activation (i.e., enhancers can be activated by stimuli) and (ii) they form a transcription memory, which is similar to dormant enhancer activation during cell differentiation [46]. The increased size of LANA NBs by the previous activation also suggested that not all of the 21 TR unit may be equally activated (tethered BRD4) in the episome. Using a strategy to sync/isolate similar classes of cell populations, we next examined the association between TR length and the magnitude of reactivation (e.g., the extent of transcription memory formation after resting for 10 days) (**Figure 5E**). Similar to the results shown above, cBu-1 TR6-KSHV-infected iSLK cells demonstrated decreased inducible gene expression per genome compared with cBu-1 TR21-KSHV-infected iSLK cells (**Figure 5F**). Decreased gene expression was accompanied by a decreased frequency or duration of BRD4 recruitment at the inducible promoter (**Figure 5G**). These results suggest that increased TR copy numbers anchor more LANA via DNA sequence-specific interactions (**Figure 3A**) and that the LANA/TR complex could harbor BRD4 near the latent episome (**Figure 3C**). An increased local concentration of BRD4 proximity to the latent episome within the 3D nucleus facilitates BRD4 delivery to the individual KSHV inducible promoter, perhaps through BRD4 interactions with transcription factors such as K-Rta (**Figure 5G**).

**Figure 5.**
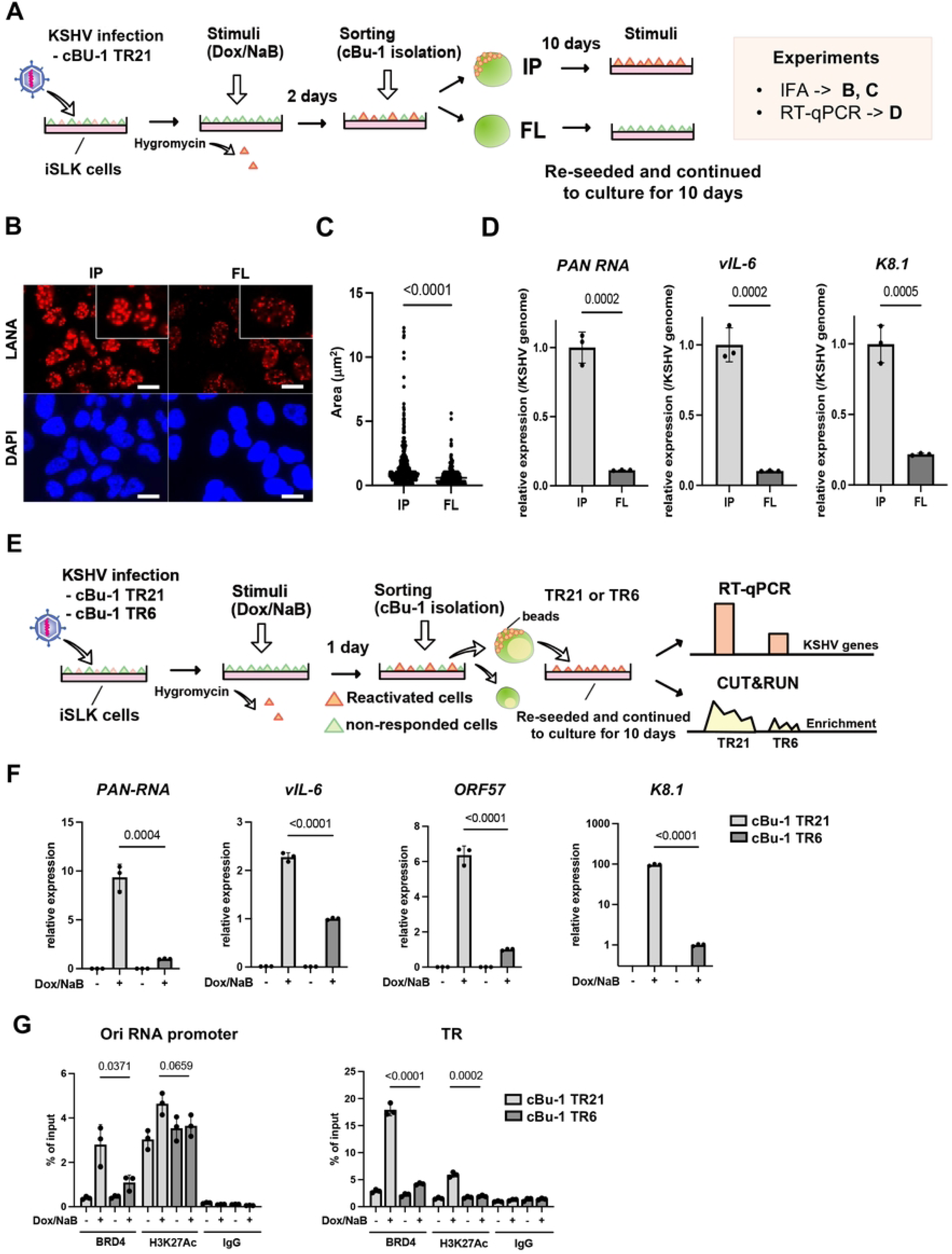
LANA NBs functions as transcription memory. **(A). Experimental workflow of the isolation of reactivated cells by cBu-1 system for the following analysis.** Two days after reactivation by sodium butyrate and doxycycline, cBu-1 expressed cells (IP) and cBu-1 non-expressed cells (FL) were sorted by magnetic beads. Cells were then cultured for ten days for the following analysis. **(B). Immunofluorescence analysis showing LANA dots in IP and FL population.** Anti-LANA rat monoclonal antibody and alexa-647 (secondary) goat anti-rat antibody were used for staining LANA in IP and FL population from cBu-1 TR21-KSHV-infected iSLK cells. Cy5 channel was used for its detection. LANA dots (upper panels) and DAPI (lower panels) are shown. Scales: 20 μm. **(C). Quantification of LANA dots.** The size of each LANA dot in IP and FL population was determined and calculated by imageJ (ver.1.53). **(D). Relative KSHV gene expression in IP and FL population.** IP and FL population were cultured for ten days, followed by reactivation by sodium butyrate and doxycycline. Two days after reactivation, total RNAs were purified to determine the viral gene expression. 18S rRNA was used as an internal standard for normalization, and relative gene expression was normalized by the viral genome. Data was analyzed using two-sided unpaired Student’s t test and shown as mean ± SD. **(E). Experimental workflow of the isolation and the comparison of reactivated cBu-1 TR21- and TR6-KSHV infected iSLK cells.** cBu-1 TR21 or cBu-1 TR6 virus were infected with iSLK cells and preapred stably infected cells with hyglomycin selection. Two days after reactivation, cBu-1 expressed cells were sorted by magnetic beads. Cells were then cultured for ten days for the following analysis. **(F). Relative KSHV gene expression in cBu-1 TR21- and cBu-1 TR6-KSHV infected iSLK cells.** cBu-1 expressed population in cBu-1 TR21- and TR6-KSHV infected iSLK cells were cultured for ten days, followed by reactivation. Re-seeded cells were reactivated again, and total RNAs were purified to determine the viral gene expression. 18S rRNA was used as an internal standard for normalization, and relative gene expression was normalized by the viral genome. Data was analyzed using two-sided unpaired Student’s t test and shown as mean ± SD. **(G). BRD4 and H3K27Ac occupancy in cBu-1 TR21- and cBu-1 TR6-KSHV-infected iSLK cells.** BRD4 or H3K27Ac enrichment of the Ori-RNA promoter and TR region was determined using CUT&RUN. Normal Rabbit IgG antibody was used as a negative control. Data was analyzed using a two-sided unpaired Student’s t-test and shown as mean ± SD.

### LANA and TR forms liquid-liquid phase-separated droplets with BRD4

BRD4 localizes at enhancers, and its intrinsically disordered regions (IDRs) form liquid-liquid phase separated (LLPS) droplets that concentrate transcription-related enzymes [16]. LANA condensates are also reported to be partially mediated by LLPS [47] and colocalize with BRD4 [5, 32, 48, 49]. Accordingly, we next examined whether LANA can form LLPS with or without BRD4 and evaluated the contribution of TR DNA fragments to LLPS formation *in vitro*. To directly visualize their interactions and droplet formation, full-length EGFP-LANA and mCherry-BRD4 were expressed with a recombinant baculovirus and purified from insect cells (**Figures 6A, upper and B, left**). As a control, we also purified EGFP or mCherry tag alone (**Figures 6A, lower and B. right**). The in vitro phase separation studies showed that EGFP-LANA did not form droplets by itself under the conditions we used (10% PEG6,000) (**Figure 6C**). However, LANA rapidly formed LLPS in the presence of mCherry-BRD4, as demonstrated by the colocalization of green and red signals in the same droplet (**Figure 6C**). The droplets were disrupted in the presence of high salt concentration or 1,6 HD (**Figure 6D**), suggesting that BRD4/LANA droplets had liquid-like properties, which is consistent with a previous report on BRD4 [16]. Notably, incubation with TR21 DNA fragments isolated from BAC16 DNA significantly increased the size of BRD4/LANA LLPS droplets, whereas a similar size of non-TR containing KSHV genome fragment isolated from the same BAC16 DNA did not (**Figures 6E and, F**). These results suggest that KSHV TR units serve as a platform for the formation of a LANA-mediated genomic enhancer. Taken together, we conclude that KSHV TR functions as a cleverly designed KSHV enhancer to maintain epigenetically active latent chromatin (**Figure 7**). Our studies support previous studies that the interaction of LANA with BRD4 is vital and central mechanism for the KSHV inducible gene regulation.

**Figure 6.**
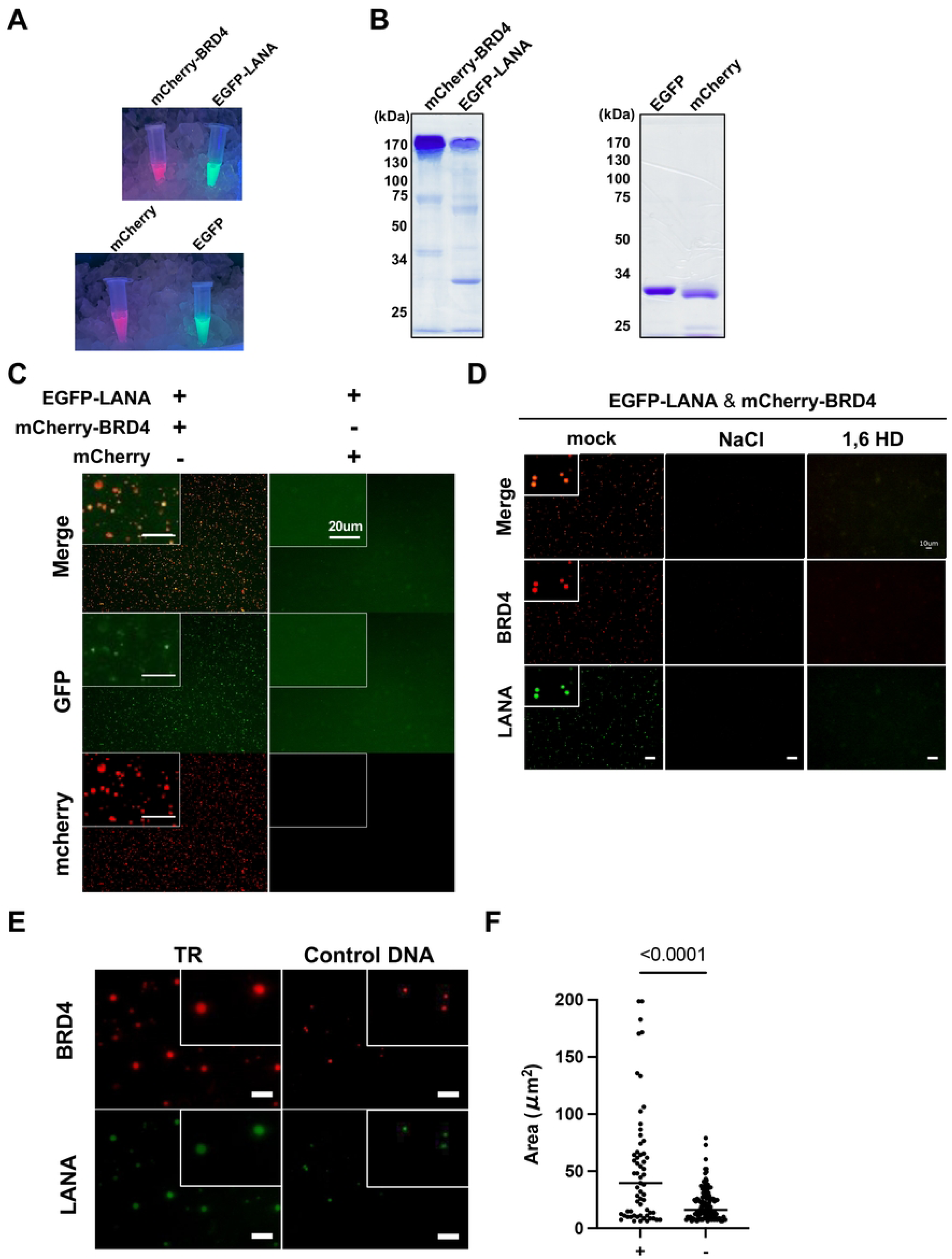
BRD4 and LANA protein form LLPS. **(A). Representative image of fluorescent proteins.** Indicated proteins were purified with Flag-beads and imaged under UV light. From left to right (up): mCherry-BRD4 and EGFP-LANA (bottom) mCherry and EGFP proteins. **(B). SDS-PAGE analysis of purified proteins**. Molecular weight markers (kDa) are indicated on the left. **(C). Representative images of LANA LLPS with or without BRD4**. EGFP-LANA (1 μM) was mixed with mCherry-BRD4 protein (1 μM) or mCherry protein (1 μM). Images were taken five minutes after the mixture of the proteins. Scales: 20 μm. **(D). Representative images of LANA and BRD4 LLPS in the presence or absence of NaCl or 1,6 HD.** EGFP-LANA (1 μM) and mCherry-BRD4 (1 μM) were mixed with sodium chloride (NaCl) (1 M) or 1,6-hexanediol (1,6HD) (10%) for ten minutes, and images were taken under fluorescence microscopy. Scales: 10 μm. **(E). Representative images of LANA and BRD4 LLPS in the presence or absence of the purified TR fragment.** EGFP-LANA (1 μM) were incubated with purified TR21 DNA fragment (50 ng) or KSHV genome lacking TR (≃ 16kbp, 50 ng) for ten minutes at room temperature. EGFP-LANA were then mixed with mCherry-BRD4 protein (1 μM) for five minutes at room temperature. Scales: 20 μm. **(F) Quantitative measurements of EGFP-LANA and mCherry-BRD4 LLPS droplet.** Droplets were determined and calculated with roundness > 0.5 by using ImageJ (ver.1.53). Data was analyzed using two-sided unpaired Student’s t test and shown as mean ± SD.

**Figure 7.**
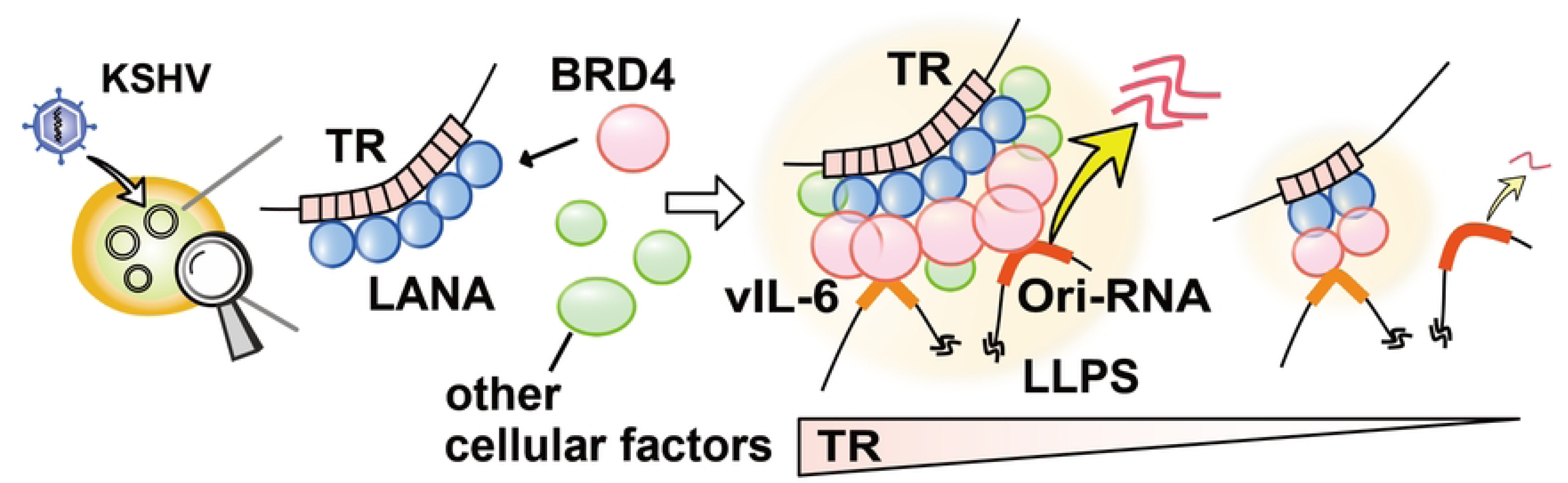
Schematic model of TR functions as an inducible enhancer for KSHV lytic gene transcription. KSHV LANA binds TR sequence specific manner and LANA recruits BRD4 with protein-protein interactions. The LANA forms LLPS in presence of BRD4, TR DNA fragment serves as a platform to recruit multiple copies of LANA to facilitate larger BRD4 containing LLPS formation. The larger LLPS (enhancer) increases a chance to interact with inducible promoters tightly packed in unique region.

## Discussion

While many key cellular transcription factors have been identified as regulators for the KSHV latency-lytic switch [39, 50-53], the mechanism by which many viral promoters are synchronously activated in response to stimuli remains less clear. Previous studies showed that TR was heavily occupied by transcription-related enzymes that are known to interact with LANA or K-Rta, and fusing TR fragments with promoters significantly increased promoter activity *in vitro*, suggesting TR could serve as a gene regulatory domain for lytic inducible promoters [5, 6]. Here, we generated recombinant KSHV to examine the significance of TR copies in KSHV replication and its mechanism of action.

To examine the association of TR enhancer activity with KSHV replication, we first established the recombinant KSHV with different TR copies and examined the effect of TR copy number on KSHV gene expression. We were hopeful to correlate strength (length) with promoter activation (lytic gene expression) by generating many recombinant KSHVs. However, our initial screening showed that the enhancer strength was not linearly associated with inducible gene expression; there may be a threshold for the TR copies necessary to trigger KSHV inducible gene expression. For example, we found that TR2 was not sufficient to reliably establish KSHV latently infected cells, and TR10 showed indistinguishable transcription profiles from TR21 in iSLK cells in reactivation. However, it is important to note that K-Rta expression is driven by an exogenous inducible cassette in the iSLK cell model, and K-Rta expression is a critical part of initiating the KSHV transcription elongation from latent chromatins. We previously showed that the K-Rta promoter region localized relatively close to TRs in 3D structure [5, 8]. Thus, the TR10 virus may have impaired inducible lytic gene expression if we use endogenous K-Rta to trigger KSHV reactivation. It is also reasonable to speculate that having a higher number of TR copies (e.g., 4.0 kb for five copies vs. 16.8 kb for 21 copies) facilitates more frequent genomic interactions with a 135-kb unique region. Consistent with the hypothesis, when we isolated vIL-6 expressing cells and compared the vIL-6 promoter with other inducible promoter activations, we found that the TR21 virus did not exhibit further recruitment of BRD4 at the vIL-6 promoter or an increase in vIL-6 expression. The results suggest that once transcription is activated, TR size has little effect. However, when we compared other inducible genes in vIL-6-expressing cells, we found significant differences in surrounding gene expression. The results clearly suggested that KSHV enhancer strength can be defined by its ability to interact with multiple inducible promoters at the same time. Tightly packed KSHV inducible promoters in the unique region may be an advantage for increasing the chance of being activated by the viral enhancer (**Figure 7**).

KSHV latent chromatin is regulated by multiple histone enzymes that modify local histones to attract specific histone-binding proteins and form topologically associated domains with cellular cohesion complexes [54]; all of these mechanisms are similar to those of cellular chromatins [55]. Accordingly, we postulated that the regulation of viral promoter activation by the viral enhancer should also be akin to host enhancer-promoter regulation. We took advantage of mini-scale chromatin and conveniently manipulated enhancer strength by reducing TR copy number with the BAC recombination system, which in turn decreased transcription factor (LANA) binding sites, thereby reducing BRD4 recruitment to enhancers and its distribution to promoters involved in transcription activation. Similarly, when we enriched the reactivatable iSLK cell population from a dish, the previous reactivated cell population demonstrated significantly higher reactivation potencies than the non-responder flow-through population. Immunofluorescence studies showed that LANA forms significantly larger dots in the highly reactive iSLK cell populations. This suggests that the contents of LANA NBs may also be dynamically regulated by the local transcription activity, which should be tightly associated with transcription memory from the previous round of transcription (signaling) activation. Future studies should investigate how tissue microenvironments, such as constitutive inflammatory signaling activation, change the protein contents at the KSHV enhancer. Related to this, we previously showed that latent chromatin with vIL-6 knockout exhibited significantly impaired reactivation potency [56], while continued external vIL-6 stimulation increased BRD4 occupancy on both viral and host chromatin, facilitating viral gene transcription and reprogramming infected monocytes [36, 56, 57]. It is exciting to speculate that the evolutionally maintaining pro-inflammatory homologs in the KSHV genome may serve to ensure viral and host cell enhancer activity, aligning with the need of the virus for replication. Further studies will be required to prove this.

Using inducible LANA degradation with auxin and BRD4 degradation with MZ, we could show that BRD4 is a central player in maintaining inducible latent chromatin. The loss of BRD4 occupancy in latent chromatin by LANA degradation also suggested that LANA plays a major role in maintaining the local concentration of BRD4 around KSHV episomes, similar to how cellular transcription factors establish cellular enhancer domains [58, 59]. However, our studies also showed that BRD4 was recruited to the inducible lytic promoters in the absence of LANA during active gene transcription, suggesting that other DNA binding proteins, such as K-Rta, are important for the BRD4 recruitment for transcription initiation. It is important to note that the Ori RNA promoter possesses K-Rta direct binding sites [5, 60].

KSHV LANA NBs are very large nuclear structures, approximately 0.2 to 5 μm with TR21. Based on LANA proximity biotin labeling, many proteins, including IFI16, BRD4 and CHD4 were present in the nuclear bodies [5, 27]. With a significantly increased local LANA concentration by specific DNA binding and its ability to form LLPS with BRD4, we speculate that large LANA NBs can capture and anchor various cellular proteins near the latent chromatin, functioning like a “Kirby” to accumulate power of transcription regulation. This is because the protein concentration of many cellular transcription factors and enzymes is below the dissociation constant for protein-protein or protein-DNA interactions when they are diffused in a large nuclear space [61]. Perhaps related to this, previous studies showed that the disruption of LANA NBs with rapid LANA protein degradation exposed the KSHV genome to cyclic GMP-AMP synthase (cGAS), leading to the degradation of the KSHV genome [35]. The cGAS is known to form a LLPS, and the formation of LLPS with DNA is a critical step for triggering cGAS-STRING pathway [62, 63]. The study suggested that cGAS is one of the IDR-containing proteins present in LANA NBs. It would be interesting to see if we could force LANA NBs to take engineered repressors or degrons to “chill” the KSHV enhancer or degrade KSHV genome, thereby preventing KSHV reactivation. Understanding the selectivity of LANA NB contents and the structure of these aggregates will therefore be important.

Since many KSHV episomes are present in a single cell, it is possible that the KSHV genomes with a specific number of TR copies or TR associated with certain protein complexes may trigger reactivation, highlighting the need for single-episomal analysis in the future. Additionally, it is of great interest to determine whether the direct repeats in other herpesviruses possess similar transcriptional regulatory mechanisms. We found that the repetitive sequences in the EBNA1 binding sites in the EBV genome vary between clinical isolates, which may be associated with the potency of the EBV reactivation.

In summary, KSHV TR functions as an enhancer for KSHV inducible promoters, and LANA-BRD4 interaction is the central mechanism for the promoter activation function. Because the activity of the enhancer was critically associated with the frequency of KSHV replication, understanding how the contents of LANA NBs are influenced by the pathogenic tissue environment or clinical drugs, should be important areas of future research.

**Table 1.**
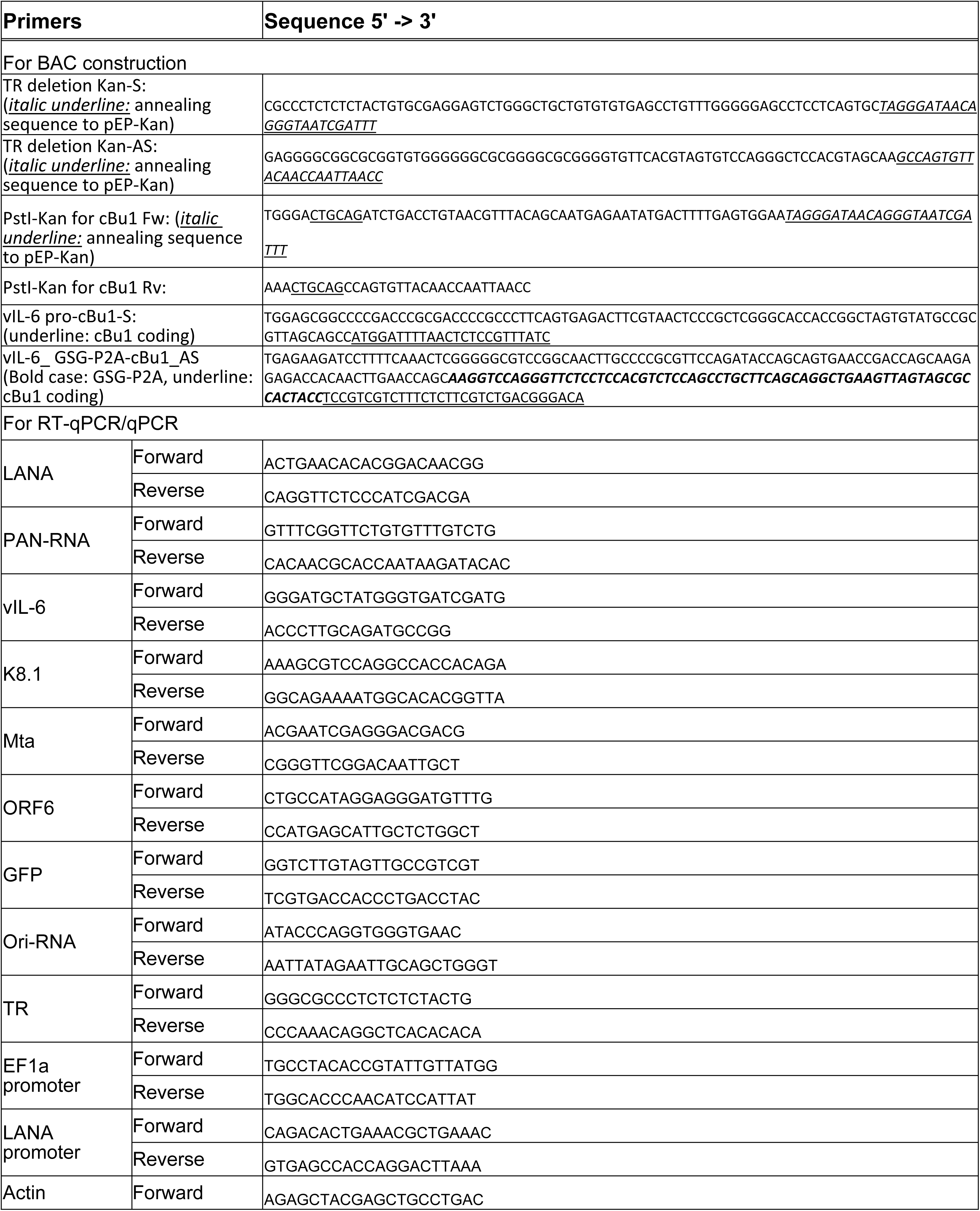
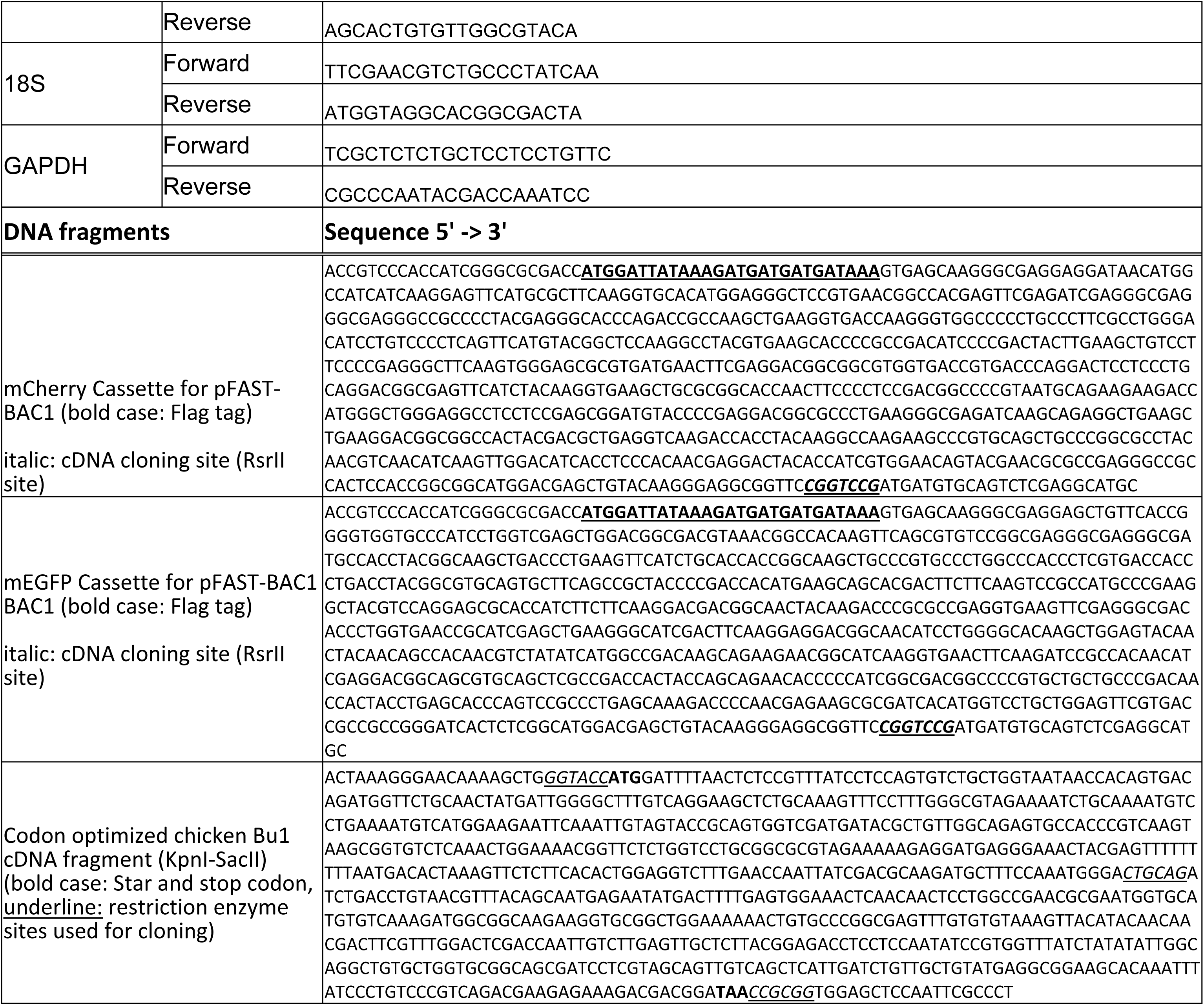
Primer and synthesized DNA fragments used in this study.

## Acknowledgments

We want to thank Dr. Robert Yarchoan (NIH/NCI) for the generous gift of the anti-vIL-6 rabbit monoclonal antibody. This research was supported by public health grants from the National Cancer Institute (R01CA290700, R21CA299587), the National Institute of Allergy and Infectious Disease (R01AI167663, R21AI186385), and American Cancer Society grant MBGI-24-1255200-01-MBG Grant DOI #: [doi.org/10.53354/ACS.MBGI-24-1255200-01-MBG.pc.gr.222219] to Y.I.

## Author Contributions

T.I., A.K., KI.N., and Y.I. designed the experiments. T.I. and K.H.W. performed flow cytometry analyses. T.I., A.K., and S.K. performed statistical analyses and visualization of the datasets. J.M.E., C.S.B.A., C.I., and Y.I. designed and prepared plasmid constructs. K.H.W., J.E., C.S.B.A., C.I., and Y.I. purified recombinant proteins. T.I. and C.S.B.A. performed LLPS formation studies. T.I. and Y.I. drafted the manuscript, and all authors edited the drafts.

## Materials and Methods

### Chemicals, reagents, and antibodies

Dulbecco’s modified minimal essential medium (DMEM), RPMI 1640 medium, fetal bovine serum (FBS), phosphate-buffered saline (PBS), Trypsin-EDTA solution, and 100 X penicillin-streptomycin–L-glutamine solution were purchased from Thermo Fisher. Puromycin and G418 solution were obtained from InvivoGen. Hygromycin B solution was purchased from Enzo Life Science. Anti-LANA rat monoclonal antibody (clone LN53) was purchased from Millipore Sigma and anti-H3K27Ac rabbit antibody (clone D5E4) was purchased from Cell Signaling Technology for immunofluorescence analysis. For proximity labeling assay, anti-LANA rabbit monoclonal antibody (clone 5L10, Millipore Sigma) and anti-BRD4 mouse antibody (E4X7E, Cell Signaling Technology) were purchased. Anti-cBu-1 mouse monoclonal antibody (biotin-conjugated) was purchased from BioVenic. Alexa 488- and 647-conjugated secondary antibody, SlowFade Gold anti-fade reagent, Lipofectamine 2000 reagent, and high-capacity cDNA reverse transcription kit were purchased from Thermo Fisher. The Quick-RNA Miniprep kit was purchased from Zymo Research, and the QIAamp DNA mini kit was purchased from QIAGEN.

### Virus preparation

Doxycycline-inducible iSLK cell lines infected with r.219 KSHV (iSLK.219 cell lines) were maintained in DMEM supplemented with 10% FBS, 1% penicillin-streptomycin-L-glutamine solution, 1μg/mL puromycin. For virus production, iSLK.219 cell lines were stimulated with 1mM sodium butylate and 1μg/ml doxycycline. Two days after chemical induction, sodium butylate, and doxycycline were removed, and cells were incubated with fresh DMEM for another 2 or 3 days. Recombinant KSHV particles were collected from the culture supernatant after centrifugation (2000rpm, 3min) and stored at -80°C until use. To quantify the encapsidated viral DNA copies, viral supernatants were first treated with DNase I (12 μg/mL) for 15 min at room temperature to degrade un-capsidated DNA. This reaction was stopped by the addition of EDTA to 5 mM, followed by heating at 70°C for 15 min. Viral genomic DNA was purified using the QIAamp DNA Mini Kit according to the manufacturer’s protocol. Elute solutions were used for real-time qPCR to determine viral copy number, as described previously [64].

### Preparation of recombinant KSHV BACs

BAC16 was a generous gift from Dr. Jae U. Jung [Cleveland Clinic Learner Center [23]]. To test the role of TR, the number of TR copies was manipulated with BAC recombination. Long primer pairs that encode a partial sequence of TR were used to prepare transfer DNA fragments for recombinant by amplifying the Kanamycin cassette from the pEP-Kan plasmid [24]. The amplified DNA fragment was transformed into BAC16 KSHV containing *E.coli* for recombination. The DNA fragments were randomly integrated into TR, which is a direct repeat of 801 bp fragments. With the I-SceI induction with L-Arabinose, we induced double-strand breaks at direct repeat regions, and subsequent induction of red recombinase with increased temperature induces random recombination within TRs. The resulting BAC clones contain different numbers of TR copies due to recombination at homologous recombination within TRs. The Kanamycin-sensitive *E.coli* clones were cultured in 10 mL of LBs, and purified BAC DNAs were digested with PstI or KpnI to estimate the number of TR copies. Based on the agarose gel electrophoresis and EtBr staining, we selected several *E.coli* clones containing different numbers of TR copies. Similarly, the GSG-P2A chicken Bu1 extracellular domain was inserted in the N-terminal region of vIL-6 (K2) as a fusion. For that, the codon-optimized cBu1 fragment was synthesized and cloned into a pBS vector. Kanamycin expression cassette with I-SceI restriction enzyme fragment with 50 bp homology arm was amplified and cloned into *Pst*I restriction enzyme site in the cBu1 fragment. Long primer pairs with homology arm with K2 N-terminal region were used to amplify the cBu1-Kan fragment, and recombination was performed by transforming purified PCR fragment into BAC containing *E.coli* as described previously. The Kanamycin cassette was removed with red-recombination, and the surrounding sequence was confirmed by sequencing. We confirmed junctions and the cBu1-vIL-6 coding sequence. Integrity of the BAC was confirmed by restriction enzyme digestion of the purified BACs. The resulting cBu-1-vIL-6 BAC was used as a backbone and generated TR deletion BACs, as described above.

### Preparation of fluorescence tagged recombinant protein

pFAST-BAC1 plasmid was first engineered to have an N-terminal Flag tag and fluorescence proteins, EGFP, mCherry, or mBFP. The restriction enzyme site, the CpoI site, was included as a cDNA cloning site to make a fusion protein with EGFP or mCherry. The cDNAs of our protein of interest were cloned into the CpoI sites, and recombinant baculoviruses were prepared with the Bac to Bac system as previously described [64, 65]. The transfer plasmid, pFAST-BAC1 vectors encoding Flag-EGFP-LANA, Flag-mBFP-LANA, Flag-mCherry-BRD4, Flag-mCherry-K-Rta, Flag-EGFP alone, Flag-mCherry, or Flag-mBFP tag alone was recombined with Bac to Bac system. Recombinant baculovirus bacmid DNA isolated from *E.coli* was transfected into Sf9 cells by using polyethylenimine (Sigma), and recombinant viruses were subsequently amplified twice. Expression of recombinant proteins was confirmed by immunoblotting with anti-Flag monoclonal antibody (Sigma). Large-scale cultures of Sf9 cells (50 ml) were infected with recombinant baculovirus at a multiplicity of infection (MOI) of 0.1 to 1.0, and cells were harvested 48 hrs after infection. Recombinant proteins were purified as described previously. The purity and amount of protein were measured by BCA assay (Invitrogen) and further confirmed by SDS-PAGE and coomassie blue staining using bovine serum albumin (BSA) as a standard.

### Immunoblotting

Cells were washed with PBS, lysed in lysis buffer (50 mM Tris-HCl [pH 6.8], 2% SDS, 10% glycerol), and boiled for 3min. The protein concentrations of the lysates were quantified with a BCA Protein Assay Kit (Thermo Fisher). Protein samples were separated by SDS-PAGE using 10% agarose gel and transferred to transfer membranes (Millipore-Sigma, St. Louis, MO, USA), which were incubated in 5% nonfat milk at room temperature for 2 hours. The membrane was incubated with the primary antibody at 4°C overnight or at room temperature for 2 hours. The membrane was then incubated with horseradish-peroxidase-conjugated secondary antibody or Alexa-647-conjugated secondary antibody at 25°C for 1 hour. For cell fractionation, cells were suspended with hypotonic buffer (20mM Tris-HCl (pH 7.4), 10mM NaCl, 3mM MgCl2, 0.5mM DTT, and proteinase inhibitor cocktail) for 15min on ice and 0.5% NP-40 was added. Cells were then centrifuged for 10 minutes at 3000 rpm to collect the supernatants (cytoplasm fraction). Pellets were washed with PBS 2 times and suspended with protein lysis buffer (nuclear fraction).

### RT-qPCR

Cells were washed with PBS, and total RNA was extracted using the Quick-RNA miniprep kit (Zymo Research, Irvine, CA, USA). Purified RNA was incubated with DNase I for 15 and reverse transcribed with the High-Capacity cDNA Reverse Transcription Kit (Thermo Fisher, Waltham, MA USA). The resulting cDNA was used for qPCR. SYBR Green Universal master mix (Bio-Rad) was used for qPCR according to the manufacturer’s instructions. Each sample was normalized to β-actin RNA, and the duct fold change method was used to calculate relative quantification. All reactions were run in triplicate using specific primers designed by PrimerQuest (Integrated DNA Technologies).

### Cleavage Under Targets and Release Using Nuclease (CUT&RUN)

CUT & RUN [66] was performed by following the online protocol developed by Dr. Henikoff’s lab with a few modifications to fit our needs. Cells were washed with PBS and wash buffer [20 mM HEPES-KOH pH 7.5, 150 mM NaCl, 0.5 mM Spermidine (Sigma, S2626), and proteinase inhibitor (Roche)]. After removing the wash buffer, cells were captured on magnetic concanavalin A (ConA) beads (Polysciences, PA, USA) in the presence of CaCl2. Beads/cells complexes were washed three times with digitonin wash buffer (0.02% digitonin, 20 mM HEPES-KOH pH 7.5, 150 mM NaCl, 0.5 mM Spermidine and 1x proteinase inhibitor), aliquoted, and incubated with specific antibodies (1:50) in 250 μL volume at 4°C overnight. After incubation, the unbound antibody was removed with digitonin wash buffer three times. Beads were then incubated with recombinant Protein A/G–Micrococcal Nuclease (pAG-MNase), which was purified from E.coli in 250 μl digitonin wash buffer at 1.0 μg/mL final concentration for 1 h at 4 °C with rotation. Unbound pAG-MNase was removed by washing with digitonin wash buffer three times. Pre-chilled digitonin wash buffer containing 2 mM CaCl2 (200 μL) was added to the beads and incubated on ice for 30 min. The pAG-MNase digestion was halted by the addition of 200 μl 2× STOP solution (340 mM NaCl, 20 mM EDTA, 4 mM EGTA, 50 μg/ml RNase A, 50 μg/ml glycogen). The beads were incubated with shaking at 37 °C for 10 min in a tube shaker at 300 rpm to release digested DNA fragments from the insoluble nuclear chromatin. The supernatant was then collected by removing the magnetic beads. DNA in the supernatant was purified using the NucleoSpin Gel & PCR kit (Takara Bio, Kusatsu, Shiga, Japan). Sequencing libraries were then prepared from 3 ng DNA with the Kapa HyperPrep Kit (Roche) according to the manufacturer’s standard protocol. Libraries were multiplex sequenced (2 × 150 bp, paired-end) on an Illumina NovaSeq 6000 system to yield ∼15 million mapped reads per sample. With separate replicated experiments, qPCR was used to examine enrichment at selected genomic regions.

### Immunofluorescence staining

Cells were cultured on coverslips and fixed with 2% PFA for 10 min at room temperature. Cells were incubated in PBS with 0.1% Triton X-100 for permeabilization for 20 min at room temperature and blocked (1% BSA, 0.05% Tween-20 in PBS) for 60-120 min at room temperature. The following primary antibodies were used: FLAG (Sigma, catalog no. F1804), H3K27ac (Abcam, catalog no. ab4729), MED1 (Abcam, catalog no. ab64965) and Alexa Fluor 647-BRD4 (Biolegend, catalog no. 683004). After staining overnight at 4 °C, cells were washed by permeabilization buffer three times. Secondary antibodies were Alexa Fluor 488 goat anti-mouse IgG (Invitrogen, 1:1,000) and Alexa Fluor 647 goat anti-rabbit IgG (Invitrogen, 1:1,000). Cells were counterstained with 4,6-diamidino-2-phenylindole (DAPI) and examined under All-in-One Fluorescence Microscope (BZ-X710, KEYENCE)

### Proximity ligation assay

The Proximity Ligation Assay (PLA) was performed by Duolink® Proximity Ligation Assay (Millipore Sigma) according to the manufacturer’s instructions. Briefly, cells were seeded on glass coverslips in a 6-well plate at a density of 3 × 10^5 cells per well and cultured overnight in DMEM medium. Cells were fixed with 2% paraformaldehyde in PBS for 10 minutes at room temperature, permeabilized with 0.1% Triton X-100 in PBS for 10 minutes, and blocked with the 1X blocking buffer in the kit. Primary antibodies, including anti-LANA rabbit antibody and anti-BRD4 rabbit antibody, were diluted (1:100) in the blocking buffer and incubated with the samples overnight at 4°C. After washing, the samples were incubated with PLA probes (anti-mouse MINUS and anti-rabbit PLUS) for 1 hour at 37°C. Ligation was performed by incubating the samples with ligase solution for 30 minutes at 37°C, followed by amplification with polymerase solution for 100 minutes at 37°C. Coverslips were mounted onto slides using 10 μl Duolink® In Situ Mounting Medium with DAPI. Images were taken by All-in-One Fluorescence Microscope (BZ-X710, KEYENCE)

### In vitro phase separation assay

To induce phase separation, purified proteins were diluted to a final concentration of 10 µM in a buffer containing 50 mM Tris-HCl (pH 7.5), 150 mM NaCl, and 1 mM DTT and 4% polyethylene glycol. The mixture was incubated at room temperature for 10 minutes. Phase separation was visually confirmed by fluorescence microscopy (BZ-X710, KEYENCE). The protein solution was loaded on a glass slide with a coverslip attached by two parallel strips of tape.

### Statistical analysis

Statistical analyses were performed using GraphPad Prism 9.4.1 software. Results are shown as mean ± SD with dots representing individual measurements. Statistical significance was determined by Student’s t-test, ratio paired t-test, or one-way ANOVA with Tukey’s multiple comparison test, and correction for false discovery rate (FDR) as described in each figure legend. FDR corrected p < 0.05 was considered statistically significant.

## Data availability statement

All data are available in the main text. Reagents will be available upon request. Reagents described in this manuscript will be available upon request.

